# Heightened Hedonic Impact and Drug Seeking Drive the Dramatic and Heritable Augmentation of Oxycodone Intake in Rats

**DOI:** 10.1101/2025.05.07.652717

**Authors:** Burt M. Sharp, Hao Chen

## Abstract

Dissecting the interplay between hedonic “liking” and motivational “wanting” is key to uncovering opioid addiction mechanisms. Using genetically diverse inbred rat strains trained on an operant oral oxycodone self-administration task, we identified “Augmenter” strains showing a surge in intake during extended (16-h) access. We analyzed 562,000 lick clusters, using a novel method of lick microstructure analysis, to distinguish oxycodone consumption from seeking. Augmented intake reflected increases in the number of both consummatory and seeking clusters across sexes. In female Augmenters, strong correlations between consumption and seeking cluster size (i.e. measure of hedonic impact) and interlick interval indicated that heightened hedonic experience during consumption directly amplified motivation to seek drug; in males, this linkage was weaker. Differentiating consummatory from seeking clusters thus provides a powerful framework for parsing hedonic and motivational components of opioid use, revealing that elevated hedonic impact drives the heritable, female-biased augmentation of oxycodone intake.

## Introduction

Opioid drugs possess a high addiction liability by engaging the brain’s reward circuitry through two distinct biopsychological processes: hedonic “liking,” the pleasure upon consumption, and motivational “wanting,” the incentive drive to seek the drug^1–3^. These components have been dissociated, revealing that the endogenous opioid system underlies the hedonic evaluation of reward (i.e., the core process of “liking”) by activating the brain’s “hedonic hotspots“^4,5^. In contrast, “wanting” is driven primarily by the mesolimbic dopamine system and the mu-opioid system^1,6^. According to incentive-sensitization theory, repeated drug exposure sensitizes the dopamine system, amplifying “wanting” into pathological craving^1^. The hallmark of addiction is compulsively “wanting” drug that persists and intensifies even as the pleasurable “liking” diminishes with chronic use^1^; hence, “wanting” and “liking” are distinct.

To measure these components, operant oral self-administration is advantageous compared to other tasks. The continuous stream of data from licking is uniquely suited to lick microstructure analysis (LMA), based on the hypothesis that the fine-grained temporal pattern of licking is a quantifiable motor output, which is dynamically modulated by neural circuits processing the sensory properties (e.g., identity, intensity) and hedonic value of a stimulus, in the context of the animal’s current physiological and motivational state. The number of licks per cluster, or cluster size (CS), correlates with the palatability and hedonic impact of a liquid reward, generating an objective measure of “liking” during consumption^7^. LMA has been applied to study ingestive behaviors, conditioned flavor preferences, and drug effects^8,9^, providing a quantitative measure of the hedonic value of a stimulus.

We developed an advanced LMA to study the reward value of oral oxycodone. We reported previously^10^ that extending self-administration sessions from 4 to 16-h revealed distinct behavioral phenotypes across genetically diverse inbred rat strains. Among strains with similar 4-h intake, extended access identified “Augmenters,” which increased consumption over 2.5-fold, and “Non-Augmenters.” We introduce a novel analysis that classifies lick clusters to separate drug consumption from subsequent seeking by examining cluster size and timing relative to reward delivery. This deconstructs operant licking into large “consumption” clusters reflecting direct reward engagement and smaller “seeking” clusters representing persistent responding to obtain the next reward. Using this framework, we test whether strain differences in intake augmentation arise from distinct changes in consumption and seeking, showing that enhanced hedonic impact drives increased seeking in female Augmenters.

## Results

### Identifying the Augmenter vs. Non-Augmenter phenotypes

We studied 52 inbred female strains (mean = 7.55 ± 0.45 rats/strain) and 47 inbred male strains (mean = 6.85 ± 0.36 rats/strain, Figure 1A). Across the entire panel, oxycodone intake (mg/kg) increased as the dose and session length increased (main effect of stage, which are unique combinations of oxycodone dose and session length: F_7,12331_ = 796.1, p < 2.0×10^−16^). Females showed higher overall intake than males (sex: F_1,745_ = 100.5, p < 2.0×10^−16^; female mean = 0.66 mg/kg, male mean = 0.26 mg/kg, average of all stages). Considering all strains, female peak intake was 1.13 ± 0.03 mg/kg in the 4-h, 0.1mg/ml stage, with a large increase to 2.03 ± 0.07 mg/kg at 16-h, while male peak 4-h intake was 0.43 ± 0.01 mg/kg, increasing to 0.74 ± 0.03 mg/kg at 16-h. Therefore, dose, session duration and sex influenced intake levels.

**Figure 1.**
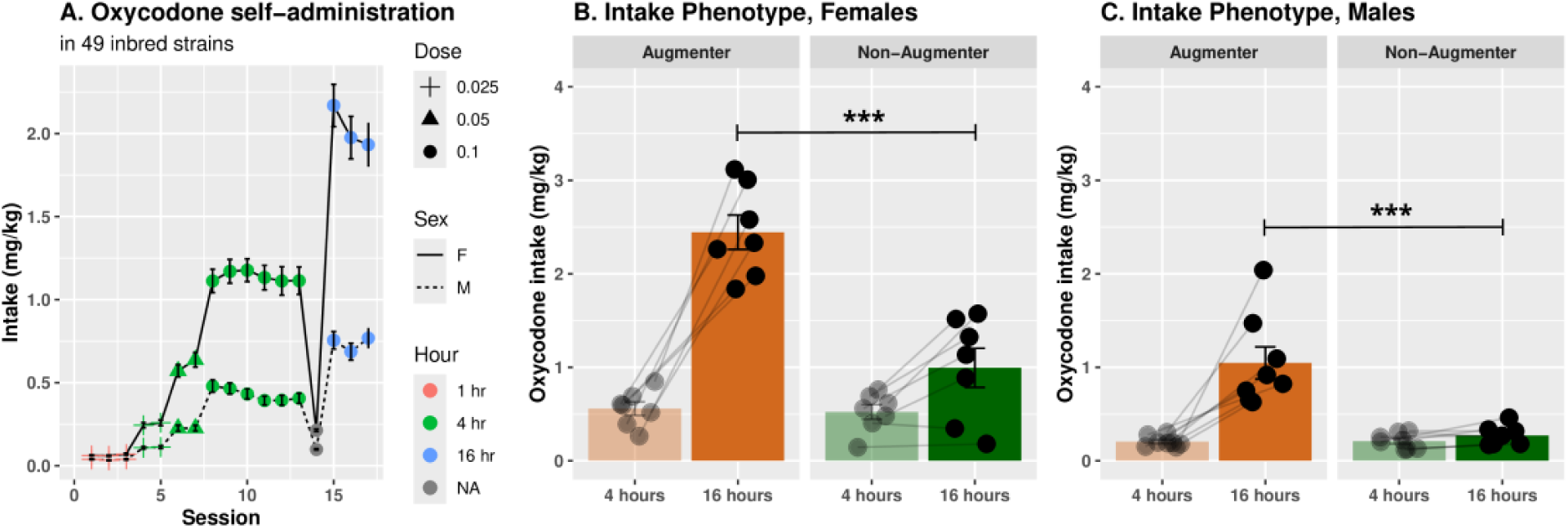
Augmenter and Non-Augmenter rat strains are distinguished by greatly increased oxycodone intake during 16-h self-administration sessions. **A.** The mean oxycodone intake in 47 male and 52 female inbred rat strains over stages of increasing drug concentration and session length. Across all strains, oxycodone intake increased significantly as self-administration dose and duration increased (p < 2.0×10^−16^), and females consumed more than males overall (p < 2.0×10^−16^). Augmenter and Non-Augmenter phenotypes are defined by the relative increase in oxycodone intake from 4-h to 16-h sessions. **B**. In female, Augmenter and Non-Augmenter strains had comparable intakes during 4-h sessions (p = 0.87). However, Augmenters consumed significantly more during the 16-h sessions, resulting in main effects of both intake phenotype (p = 0.001) and session duration (4h vs. 16h; p < 5.4×10^−07^). **C**. In males, intake did not differ between phenotypes in 4-hour sessions (p = 0.96), but Augmenters consumed more in the 16-h sessions. This was supported by main effects of intake phenotype (p < 0.001) and session duration (p < 0.001). ***: p < 0.001

Rat were divided into two *intake phenotypes*, Augmenters (7F and 8M strains) and Non-Augmenters (7F and 8M strains), based on their increase in oxycodone intake from the 4-h, 0.1mg/ml to the 16-h, 0.1mg/ml stages (see Methods). In females (Figure 1B), we found main effects of intake phenotype (F_1,12_ = 18.8, p = 0.001) and session length (4-h vs 16-h; F_1,12_ = 92.8, p < 5.4×10^−7^), along with an interaction of intake phenotype x session length (F_1,12_ = 33.3, p = 8.9×10^−5^). In females at 4-h, mean intakes were similar in Non-Augmenters and Augmenters (0.52 ± 0.08 vs 0.56 ± 0.07mg/kg, respectively; p = 0.87); however, at 16-h, Augmenters demonstrated much higher intake than Non-Augmenters (2.44 ± 0.18 vs. 0.99 ± 0.21 mg/kg, respectively; p < 0.0001). Similarly, in males (Figure 1C), main effects were intake phenotype (F_1,14_ = 18.1, p <0.001) and session length (F_1,14_ = 26.1, p <0.001), as well as an intake phenotype x schedule interaction (F_1,14_ = 19.5, p < 0.001). Similar to females at 4-h, male Non-Augmenter oxycodone intake did not differ from Augmenters (0.21 ± 0.03 vs 0.21 ± 0.02 mg/kg, respectively; p = 0.96), but 16-h intake was much greater in Augmenter males than Non-Augmenters (1.05 ± 0.17 vs 0.27 ± 0.04 mg/kg, respectively; P < 0.001). Therefore, in both sexes, augmented oxycodone intake was detected only in the Augmenters during 16-h sessions.

### Analysis of total active and inactive licks at each stage of oxycodone self-administration

Analysis of total active spout licks in female Augmenter and Non-Augmenters (Figure 2A) showed main effects of intake phenotype (F_1,12_ = 8.73, p = 0.01) and stage (F_3,36_ = 31.7, p < 0.001), and an intake phenotype x stage interaction (F_3,36_ = 13.59, p < 0.001). There was no difference in active licks between female Augmenters and Non-Augmenters during the 4-h stages (0.025mg/ml, 4-h: p = 0.06; 0.05mg/ml, 4-h: p = 0.09; 0.1mg/ml, 4-h: p = 0.32). However, Augmenters licked far more than Non-Augmenters at 16-h (2512.5 ± 200.8 vs 619.9 ± 147.9, respectively; p < 0.001). For female inactive spout licking (Figure 2A), there was a main effect of stage (F_3,36_ = 8.81, p < 0.001), but not of intake phenotype (F_1,16_ = 2.76, p = 0.12) or intake phenotype x stage (F_3,36_ = 1.45, p = 0.24).

**Figure 2.**
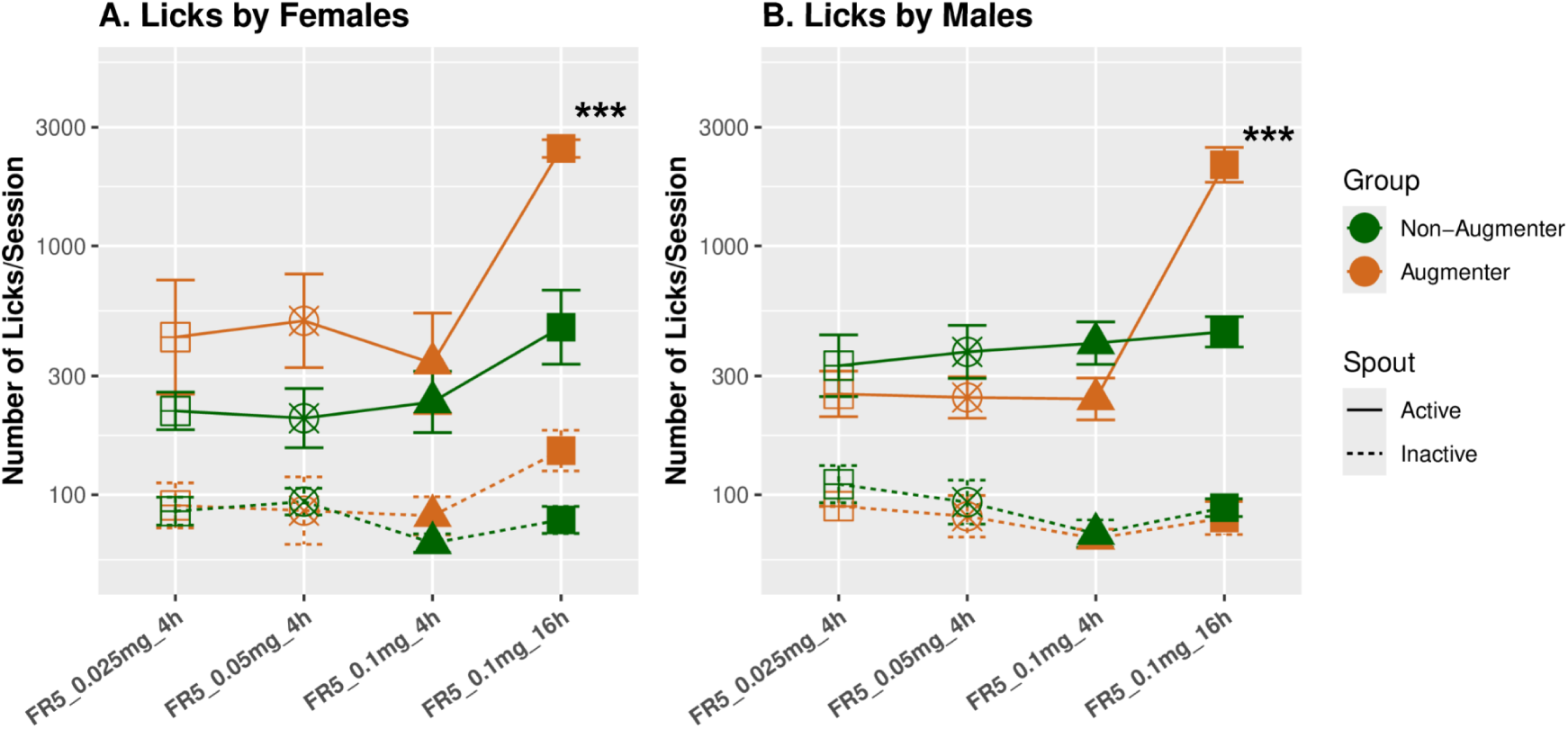
Phenotypic differences in licking patterns and microstructure during oxycodone self-administration. Total licks on the active and inactive spouts for Augmenter and Non-Augmenter rats across stages of self-administration in females (A) and males (B). For active spout licks, a significant interaction between self-administration stage and intake phenotype was found in both females (p = 0.001) and males (p < 0.001). Specifically, Augmenters licked more than Non-Augmenters at 16-h (p < 0.001, for both sexes). Licks on the inactive spout did not differ between phenotypes at any stage, demonstrating consistent spout discrimination. *** p < 0.001 between Augmenter and Non-Augmenter groups.

Similarly, in males, active spout licking (Figure 2B) showed main effects of intake phenotype (F_1,14_ = 5.8, p = 0.03) and stage (F_3,42_ = 33.8, p < 0.001), as well as an intake phenotype x stage interaction (F_3,42_ = 10.12, p < 0.001). There was no difference in active spout licking between Augmenters and Non-Augmenters in 4-h stages (0.025mg, p = 0.54; 0.05mg, p = 0.34; 0.1mg, p = 0.35). However, Augmenter males had far more active spout licks than Non-Augmenters at 16-h (2300.0 ± 327.8 vs 484.1 ± 68.1; p < 0.001). Inactive licking (Figure 2B) showed no main effects of intake phenotype (F_1,14_ = 1.60, p = 0.23) or stage (F_3,42_ = 1.18, p = 0.16), Therefore, only in 16-h self-administration sessions did male Augmenters emit a significantly greater number of active spout licks than Non-Augmenters, and inactive licks were unaffected

### Lick cluster size (CS) is more responsive to oxycodone oral self-administration than interlick interval (ILI)

CS increased while ILI decreased across self-administration stages of all rats (CS: F_4,424_ = 35.0, p < 2.2×10⁻¹⁶; ILI: F_4,334_ = 12.85, p < 0.001, Figure 3A and 3B). Post-hoc tests confirmed that the 16-h stage differed from all others, exhibiting the largest CS and shortest ILIs (ps < 0.001). CS and ILI were strongly negatively correlated on the active spout (r = -0.445, p < 2.2×10⁻¹⁶), indicating progressively faster and more persistent licking over time. We calculated the incremental change of CS and ILI from the first 4-h stage (0.025 mg/ml) to the 16-h stage in active spout and then normalized to its first 4-h stage. The mean change in the CS (0.40 ± 0.08 SE) was greater (p < 0.001) than in ILI (-0.03 ± 0.01 SE). The effect size for the CS analyzed by Cohen’s d from the initial 4-h to 16-h was also larger than that of ILI (CS: d = 0.93, large effect; ILI: d = -0.32, small effect). Therefore, the primary change in lick microstructure over time was an increase in the number of licks per cluster, whereas altered timing between individual licks was comparatively modest. Hence, we focused subsequent analyses of consumption vs seeking on CS.

**Figure 3.**
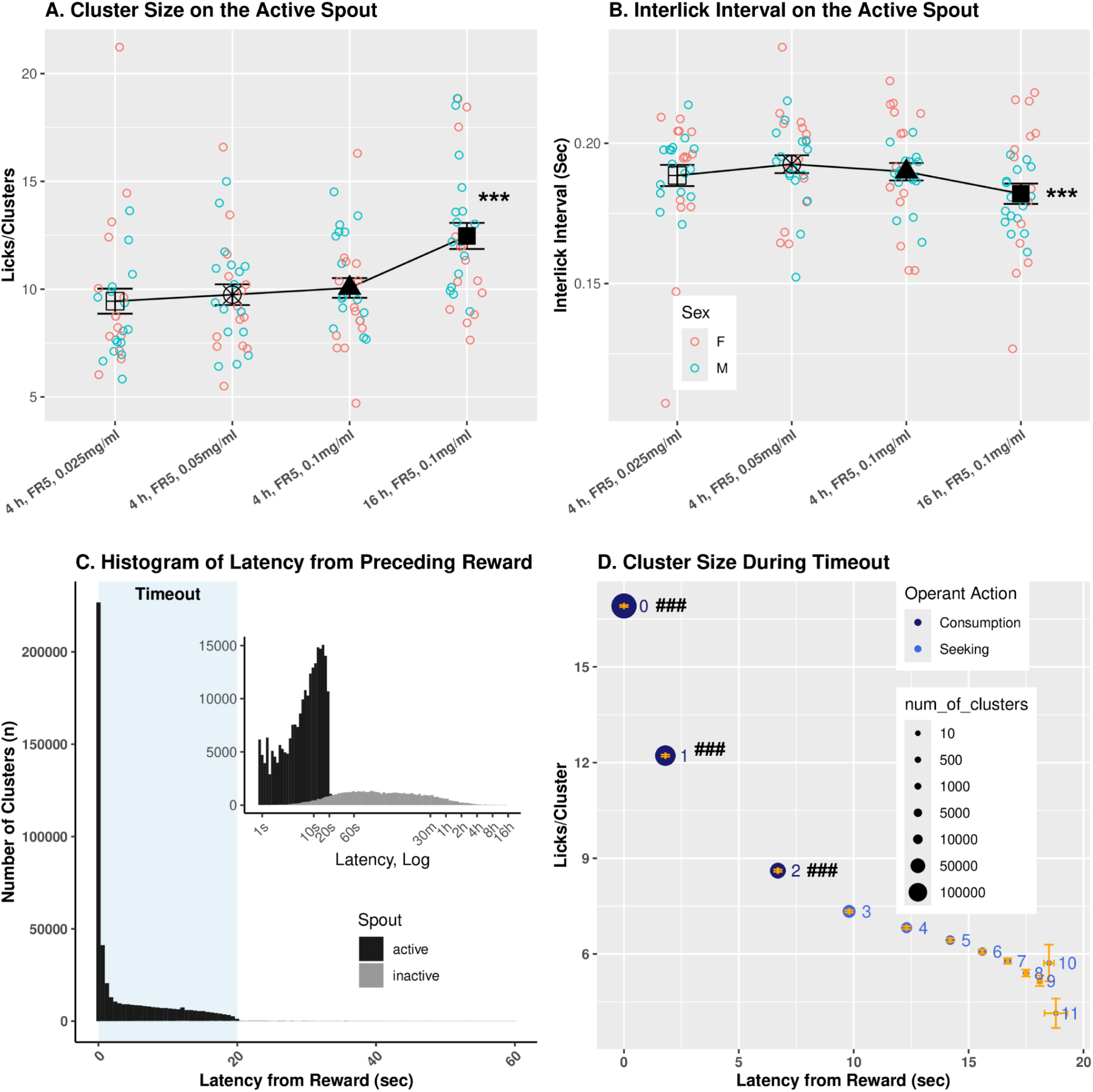
Classification of Active Spout Licking Behavior into Consummatory vs. Seeking Lick Clusters. Rats received an oral oxycodone reward after completing a sequence of five licks. Each reward was immediately followed by a 20-second “timeout” period, during which no rewards were delivered. **A.** Change in CS across four stages of self-administration. **B.** Changes in ILI across four stages of self Administration. The dynamic range was significantly greater for CS than for ILI (p < 0.001), indicating that rats primarily increased their intake by taking larger bursts of licks rather than reduce the time between individual licks. **C.** *A histogram of the latency of all lick clusters generated by all rats (Table 1).* Latency is the time from a reward to the first lick of each subsequent cluster, including clusters during or after the timeout, up to the next reward.As expected, the vast majority (98.1%) of all lick clusters on the active spout (black) occurred within the timeout period. The inset extends the data to the entire (4-h or 16-h session) using a log-transformed time axis. Licks on the inactive spout (grey) are shown for comparison. **D.** *Operational definition of consummatory and seeking lick clusters.* Mean lick CS of clusters 1-11 is plotted as a function of their sequential positions following reward delivery (n = 226,752 rewards). A significant decay in CS, reflecting a decline in response vigor, was observed for the two clusters following the reward-triggering cluster (position 0). This dynamic is characteristic of a consummatory period, as response vigor naturally declines during ingestion of the fixed-volume (60 µL) reward until it is fully consumed. Based on these characteristics, clusters at positions 0-2, which declined sharply in size, were classified as consummatory, whereas those from position 3 onward, where CSs declined gradually, were classified as seeking. Following consumption, the persistent licking for an unavailable reinforcer during the timeout is interpreted as a measure of motivational drive. The size of each data point is proportional to the number of clusters observed at that position. ***: p < 0.001, compared to other stages; ###: p < 0.001, comparing cluster size to the subsequent cluster.

### Classifying the function of licks as either Consummatory or operant drug seeking

Figure 3A shows the frequency of all lick clusters (n = 562,210, for all 4-h and 16-h sessions, Table 1) emitted by all rats in the study at increasing latency from the preceding reward. The vast majority (98.1%) of all lick clusters on the active spout occurred within the 20-second timeout period. This is expected because the five cumulative licks emitted *after* a timeout will trigger a new reward, thereby becoming a “reward-triggering cluster” that initiates a new timeout period. In contrast, only 10.1% of clusters on the inactive spout occurred within the timeout period, most likely due to rats sampling a different spout after receiving a reward.

**Table 1.**
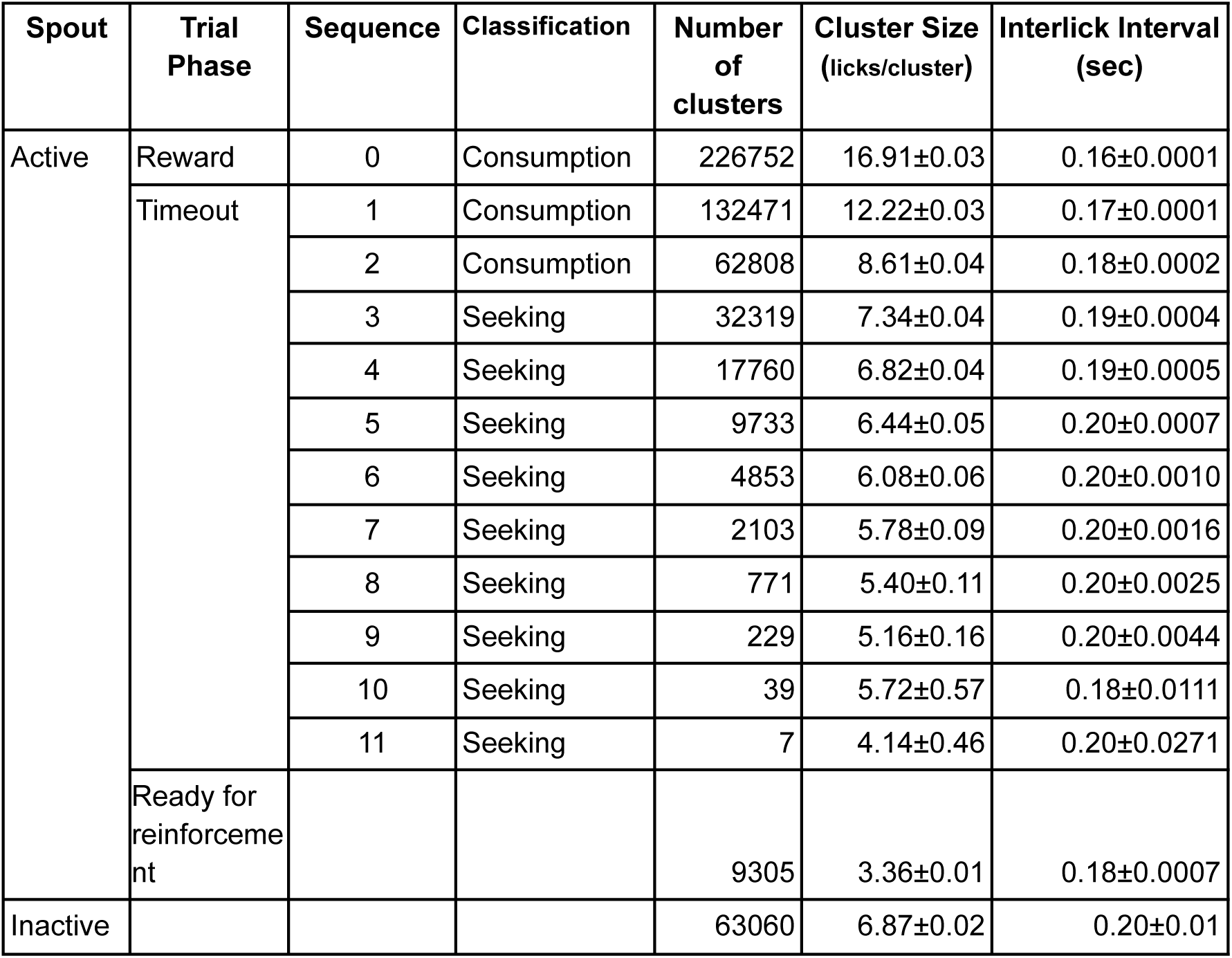
Number of clusters and their classifications.

We analyzed the microstructure of licking on the active spout to differentiate consummatory from seeking behaviors within the timeout following oxycodone reward. Our dataset contained 226,752 rewards, each initiated a 20-second timeout period (Table 1). We grouped all 489,845 clusters that occurred during these timeout periods by their sequential positions relative to the reward-triggering cluster (cluster 0, Figure 3D). CS significantly decreased as a function of the sequential position of each cluster (F_11,489054_ = 1129.3, p < 2.2×10^−16^). Specifically, the reward-triggering clusters (i.e., cluster 0, 16.91±0.03 licks/cluster) was significantly larger (p < 0.0001) than the subsequent cluster (i.e., cluster 1; 12.22 ± 0.03 licks/cluster), which was larger (p < 0.0001) than the next cluster (i.e. cluster 2; 8.61 ± 0.04 licks/cluster). CS further decreased (p < 0.0001) between the 2nd and 3rd clusters, but the size of the 3rd cluster was not different (p = 0.40) from the 4th cluster. All subsequent comparisons of CS between adjacent clusters were not significant (p > 0.05). Hence, the reward-triggering cluster and the two lick clusters immediately thereafter (i.e., 1 and 2) were significantly larger (i.e. contained more licks) than all subsequent lick clusters.

The initial sharp decline of CS is characteristic of a consummatory period during which response vigor naturally declines throughout the process of ingesting the fixed-volume (60 µL) reward. We, therefore, classify the larger early clusters (i.e., 0-2), which occur within approximately 8.5 sec after the reward (Figure 3B), as “Consumption” clusters. In contrast, the smaller clusters (i.e., 3-11), occurring later in the timeout period and with a more gradual change in CS, are classified as “Seeking” clusters. A progressive decay in cluster size was associated with a decrease in the number of lick clusters emitted (represented by dot size in Figure 3B) as a function of increasing post-reward latency. Seeking clusters, which represent the sustained licking emitted to obtain more drug when it is temporarily unavailable, are interpreted as a measure of motivation. Notably, intake phenotype did not influence the sequential changes in CS (F_2,14352_ = 0.99, p = 0.37). Under the fixed-ratio 5 schedule, all clusters outside the timeout interval are capped at 4 licks and are thus excluded from further analysis. As shown in grey in Figure 3A, licking on the inactive spout was minimal. This affirms the functional specificity of the seeking clusters generated during the timeout.

In alignment with CS, ILI increased as a function of sequential position within the timeout period (F_11,473958_ = 303.2, p < 2.2×10^−16^, Table 1). Post-hoc analysis found ILI increased significantly between the first three adjacent clusters (ps < 0.001) and remained stable between subsequent clusters (p < 0.05, Table 1).

### The frequency of lick clusters during consumption vs seeking in Augmenters and Non-Augmenters

We next compared the number of consumption and seeking clusters at the active spout in Augmenter and Non-Augmenter strains by experimental stage, sex, and intake phenotype (Figure 4). To mitigate bias from minor differences in sample size across strains, data were first averaged on a within-subject basis across all test sessions conducted within each stage (shown in the x-axis of Figure 4). These individual averages were then used to compute a final mean for each sex-specific strain. Inclusive analysis of the four stages showed more consumption clusters (39.6 ± 3.5) than seeking clusters (10.0 ± 1.3; F_1,183_ = 196.7, p < 2.2×10^−16^) per self-administration session, in agreement with Figure 3B and Table 1. We identified main effects of stage (F_3,183_ = 61.1, p < 2.2×10^−16^) and intake phenotype (Augmenters > Non-Augmenters; F_1,32_ = 23.3, p < 0.001). The effect of sex was not significant (F_1,79_ = 1.7, p = 0.2). Additionally, an interaction between stage × intake phenotype emerged (F_3,183_ = 35.7, p < 2.2×10^−16^).

**Figure 4.**
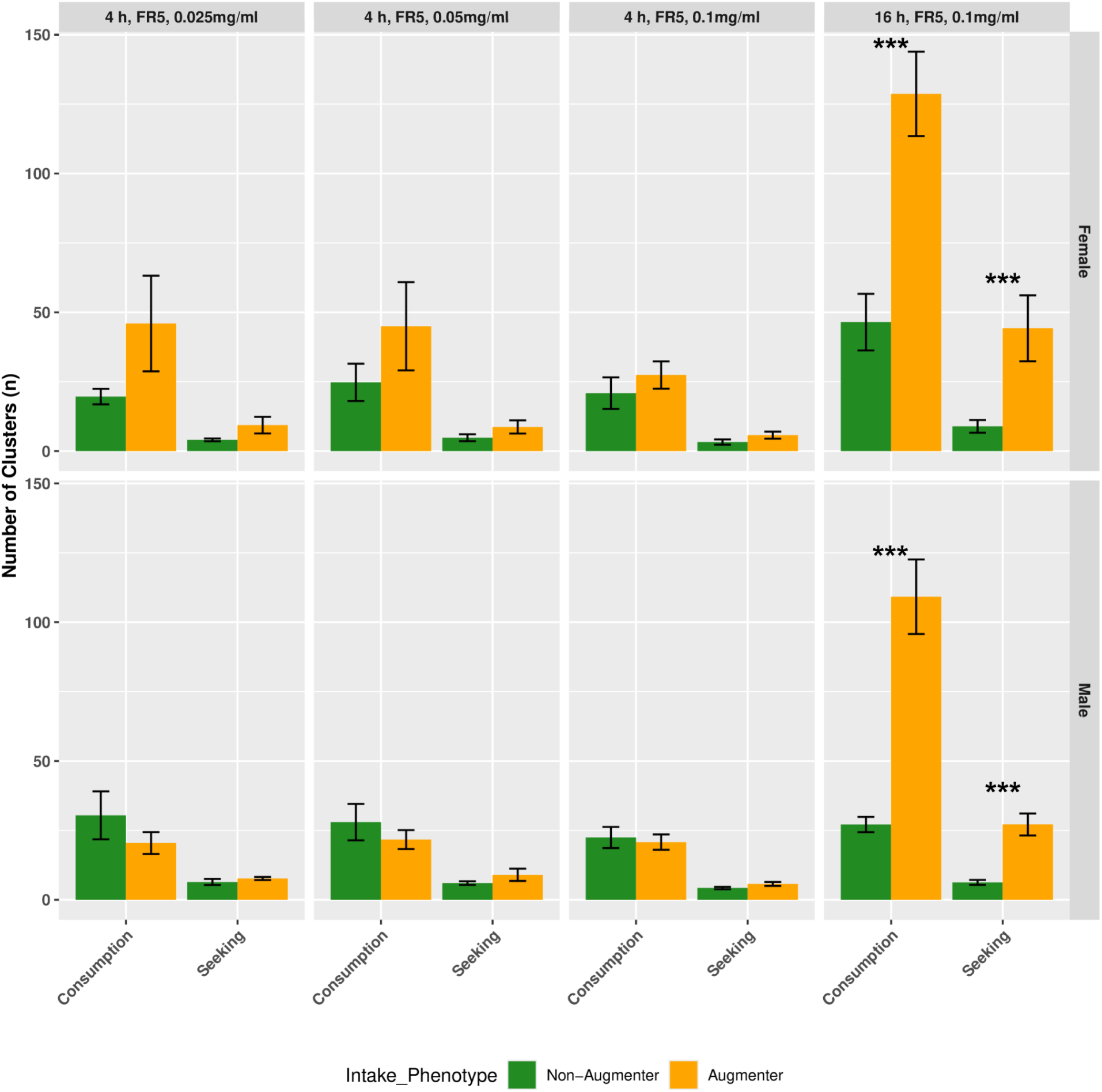
Augmenter rats exhibit more consummatory and seeking lick clusters during 16-hour self-administration (SA) sessions. The panels show the number of lick clusters on the active spout for Augmenter and Non-Augmenter rats by sex [Females (F, top); Males (M, bottom)] and cluster type [(left) Consummatory; (right) Seeking]. Overall analysis revealed main effects for all factors: self-administration Stage, number of clusters varied across stages (p < 2.2×10^−16^); Sex, females emitted a greater number of clusters than males (p = 0.031); Intake Phenotype, Augmenters emitted more clusters than Non-Augmenters (p = 6.4e-05). Crucially, while cluster counts were similar between phenotypes during the standard 4-hour sessions, Augmenters of both sexes exhibited a pronounced increase in the frequency of both consummatory and seeking clusters during the three 16-hour extended access sessions (p < 0.0001 for all comparisons). Data are mean ± SEM for all sessions at each stage.

The influence of intake phenotype on the number of clusters became progressively more pronounced across experimental stages. Specifically, at the 4-h 0.025mg/ml stage, the effect was not significant (F_1,26_ = 1.36, p = 0.25), nor at the 4-h 0.05mg/ml stage (F_1,26_ = 1.19, p = 0.28). However, a trend emerged at the 4-h 0.1mg/ml stage (F_1,27_ = 4.11, p = 0.05), and was robust by the final 16-h (F_1,19_ = 12.42, p < 0.001). Post-hoc tests revealed no difference in the number of consumption or seeking clusters between Augmenters and Non-Augmenters during the 4-h, 0.1mg/ml stage in either sex (all ps > 0.3). However, during the 16-h stage, Augmenters emitted far more consumption clusters than Non-Augmenters in both females (Aug, 128.7 ± 15.2 vs Non-Aug, 46.5 ± 10.2; p < 0.0001) and males (Aug, 109.2 ± 13.4 vs Non-Aug, 27.1 ± 2.8; p < 0.0001).

Similarly, Augmenters emitted more seeking clusters than Non-Augmenters during the 16-h stage in both females (Aug, 44.2± 11.9 vs Non-Aug, 8.9 ± 2.3; p < 0.0001) and males (Aug, 27.1 ± 4.9 vs Non-Aug, 6.2 ± 0.9; p < 0.0001). Hence, the enhanced intake observed in Augmenters during extended 16-h self-administration sessions is driven by an increased frequency of both consumption- and seeking-related licking bouts.

### Consumption and seeking cluster size reveal heritable contributions to oxycodone reward and motivation

We analyzed the average size of consumption and seeking clusters in Augmenter and Non-Augmenters (Figure 5). Across both phenotypes, consumption clusters (11.56 ± 0.44 licks/cluster) were larger than seeking clusters (7.41 ± 0.23 licks/cluster; F_1,180_ = 106.6, p < 2.2×10^−16^). Significant main effects were found for self-administration stage (F_3,180_ = 3.82, p = 0.01), with CS increasing at the 16-h stage; sex (F_1,87_ = 4.7, p = 0.03), with males (9.65 ± 0.29 licks/cluster) had larger clusters than females (9.29 ± 0.50 licks/cluster); and intake phenotype (F_1,31_ = 31.9, p = 0.01), with augmenters showed larger clusters.

**Figure 5.**
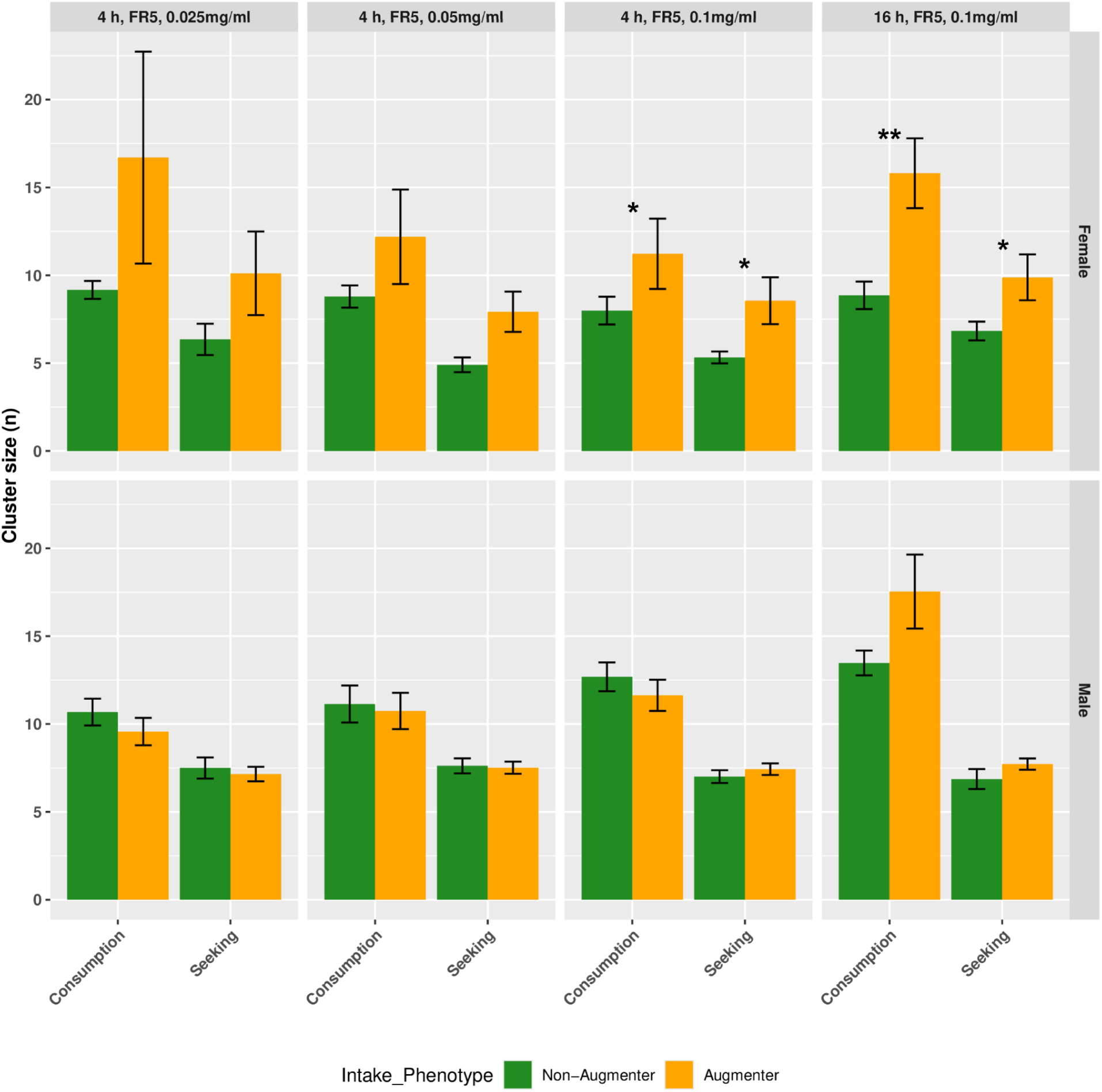
Female augmenter rats exhibit larger consummatory and seeking lick clusters during 4-h and 16-h self-administration sessions. Overall, consumption clusters were consistently much larger (i.e.,licks/cluster) than seeking clusters (p < 2.2×10^−16^). Combined analysis of all 6 panels found main effects of stage (p = 0.01), with CS generally increasing in the 16-h stage; sex (p = 0.03), with male CS greater than female; and intake phenotype (p = 0.01), with Augmenters having slightly larger CS than Non-Augmenters. CS generally increased during extended (16-h) access compared to prior sessions and differed slightly between sexes and phenotypes. At both 4-h and 16-h with 0.1mg/ml, an increase in both consumption and seeking CS was observed in female Augmenters vs Non-Augmenters (p < 0.05). Data are mean ± SEM.

The influence of intake phenotype on CS also became progressively more pronounced across self-administration stages. At the 4-h 0.025 and 0.5mg/ml stages, the effect was not significant (F_1,26_ = 2.63, p = 0.12; F_1,26_ = 3.26, p = 0.08, respectively). However, a trend emerged at the 4-h 0.1mg/ml stage (F_1,23_ = 4.11, p = 0.05), and the effect became robust by the 16-h stage (F_1,29_ = 13.4, p < 0.001). Similarly, the main effect of sex became significant only at the 0.1mg/ml stages (4-h: F_1,46_ = 5.3, p = 0.03); 16-h: F_1,50_ = 4.4, p = 0.04). There was also significant interaction between sex and intake phenotype at the 4-h 0.1mg/ml stage (F_1,41_ = 5.38, p = 0.03).

Post-hoc analysis found that female Augmenters had larger consumption CS than Non-Augmenters at the 4-h 0.1mg/ml (p = 0.03) and 16-h stages (p = 0.001). Female Augmenters also had larger CS than Non-Augmenters in both 4-h 0.1mg/ml (p = 0.03) and 16-h stages (p = 0.04). In male, we found a trend (p = 0.06) for the consumption clusters at the 16-h stage. These findings suggest that the perceived appetitive value of oxycodone and motivation to obtain oxycodone contribute to phenotype differences, especially in females.

Heritability (h^2^) of CS during consumption vs seeking phases was: females, during consumption, 0.34 and 0.29 at 4-h 0.1mg/ml and 16-h, respectively, and during seeking, 0.04 and 0.11,respectively; males, during consumption, 0.23 and 0.08, respectively, and during seeking, 0.14 and 0.11. These h^2^ values are consistent with heritability of the enhanced hedonic value of oxycodone during consumption in female Augmenters.

**Supplemental Figure 1.**
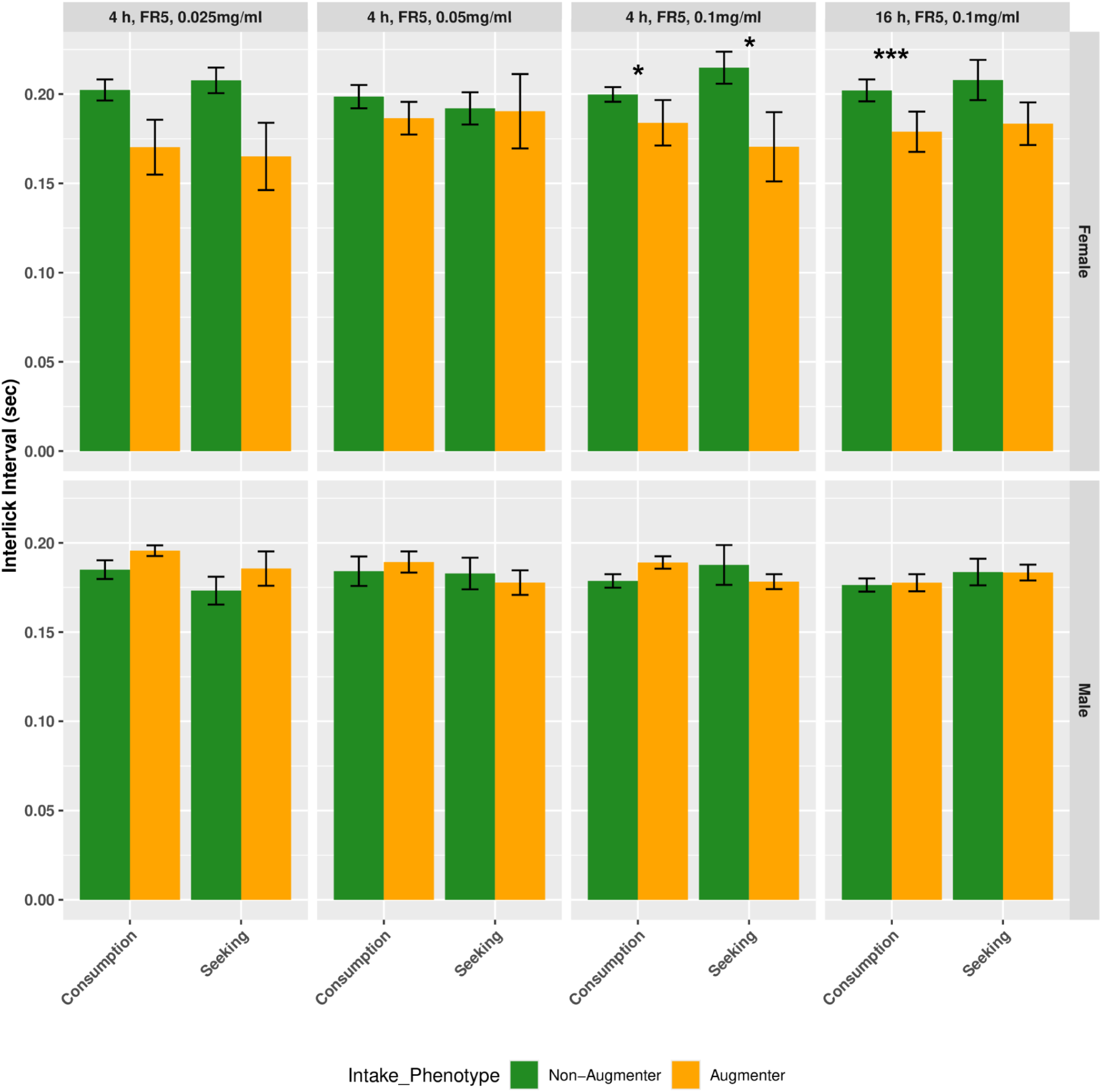
Female augmenter rats exhibit smaller consummatory and seeking interlick intervals during 4-h and 16-h self-administration sessions. Across all four self-administration stages, there were no significant differences in the interlick intervals in consumption vs seeking clusters or between Augmenters and Non-augmenters. Although males generally displayed faster licking (shorter ILI) than females, no significant overall differences in ILI were found between Augmenter and Non-Augmenter phenotypes for either cluster type. However, the main effects of sex (p < 0.001) was significant, with longer ILIs (i.e. slower licking) in females than males. In females, Augmenters exhibited shorter consumption and seeking ILIs than Non-augmenters at the 4-h, FR5, 0.1mg/ml stage. At the 16-h stage, the difference between phenotypes was large for consumption (p = 0.0001) and showed a trend for seeking (p = 0.055). Data are mean ± SEM.

### Female augmenters show shorter ILI in both consumption and seeking clusters

ILI of consumption and seeking clusters across the dataset is shown in Supplementary Figure 1. Overall, males had slightly shorter ILIs than females (0.178 ± 0.001 s vs 0.181 ± 0.003 s; F_1,181_ = 36.3, p < 0.001), with no significant overall differences between consumption and seeking clusters (0.18 ± 0.002 vs 0.18 ± 0.003 s; F_1,171_ = 0.19, p = 0.67) or between Augmenters and Non-augmenters (F_1,80_ = 0.20, p = 0.67). A trend for a three-way interaction among sex, intake phenotype, and consumption vs seeking was observed (F_3,170_ = 2.27, p = 0.07). Focusing on females, Augmenters exhibited shorter ILIs than Non-augmenters at the 4-h, FR5, 0.1mg/ml stage, consumption ILIs were 0.174 ± 0.012 s vs 0.192 ± 0.004 s (p = 0.028), and seeking ILIs were 0.162 ± 0.018 s vs 0.200 ± 0.006 s (p = 0.029). At the 16-h stage, the difference was larger for consumption (0.171 ± 0.010 s vs 0.189 ± 0.005 s; p = 0.0001) and showed a trend for seeking (0.175 ± 0.010 s vs 0.198 ± 0.012 s; p = 0.055). These findings indicate that in females, Augmenters selectively modulate licking output during both consumption and seeking, and differences between intake phenotypes become more pronounced with extended drug exposure.

### Consumption and Seeking Behaviors are Tightly Linked in Female Augmenters

To examine how consumption and seeking behaviors are coordinated, we analyzed correlations between lick microstructure parameters by sex and intake phenotype. In female (Figure 6) Augmenters at the 4-h stage, consumption and seeking ILIs were strongly positively correlated (R = 0.92, p = 0.003; panel A4), and consumption ILI was negatively correlated with seeking CS (R = -0.95, p = 0.001; panel A3). Thus, faster consumption licking was associated with larger and faster seeking clusters at 4-h. No significant correlations were observed in female Non-Augmenters during 4-h sessions. During 16-h sessions, lick microstructure in female Augmenters became more tightly coordinated. The positive correlation between consumption and seeking ILIs persisted (R = 0.98, p = 0.001; panel A4), and faster consumption continued to predict larger seeking clusters (R = -0.79, p = 0.04; panel A3). In addition, larger consumption clusters were now strongly associated with larger seeking clusters (R = 0.78, p = 0.04; panel A1) and faster seeking (seeking ILI: R = -0.82, p = 0.02; panel A2), indicating that both directional relationships were fully expressed during extended access. Therefore, well coordinated fast and persistent licking during the consumption phase led to more vigorous drug seeking immediately following consumption in female augmenters during the 16-h stage. In contrast, female Non-Augmenters displayed only a positive correlation between consumption and seeking ILIs at 16-h (R = 0.85, p = 0.03; panel B4), showing that only limited temporal coordination emerges under extended access.

**Figure 6.**
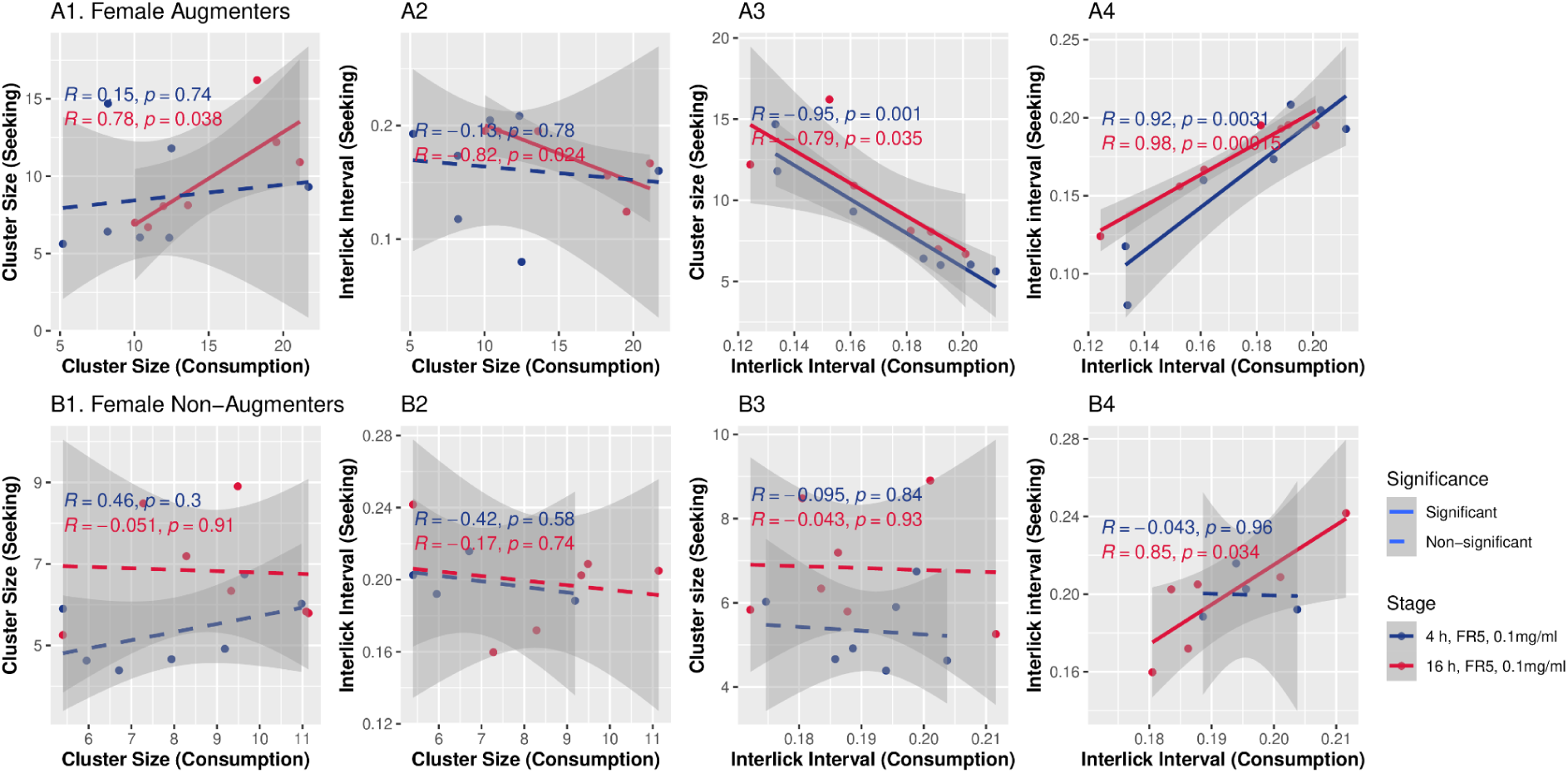
Correlations between lick microstructure parameters during consumption and seeking in female rats. The relationships between CS and ILI during consumption and seeking in female Augmenters (top row, A) and Non-Augmenters (bottom row, B). From left to right, the graphs display the correlation between: (1) consumption CS and seeking CS, (2) consumption CS and seeking ILI, (3) consumption ILI and seeking CS, and (4) consumption ILI and seeking ILI. Each point represents a strain mean. Colors define the 4-hour (blue) and 16-hour (red) sessions. The best fit line is solid for significant (p < 0.05) and dotted for non-significant correlations.

In males (Figure 7), no correlation was observed between consumption and seeking parameters at the 4-h stage in either intake phenotype. At the 16-h stage, consumption ILI was significantly correlated with seeking ILI in both Augmenters (R = 0.77, p = 0.02; panel A4) and Non-augmenters (R = 0.86, p = 0.006; panel B4). Notably, as observed in females, consumption CS predicted seeking CS in Augmenters (R = 0.75, p = 0.03; panel A1), indicating that this coupling of consumption and seeking cluster structure is a unique feature of the Augmenter phenotype shared across sexes.

**Figure 7.**
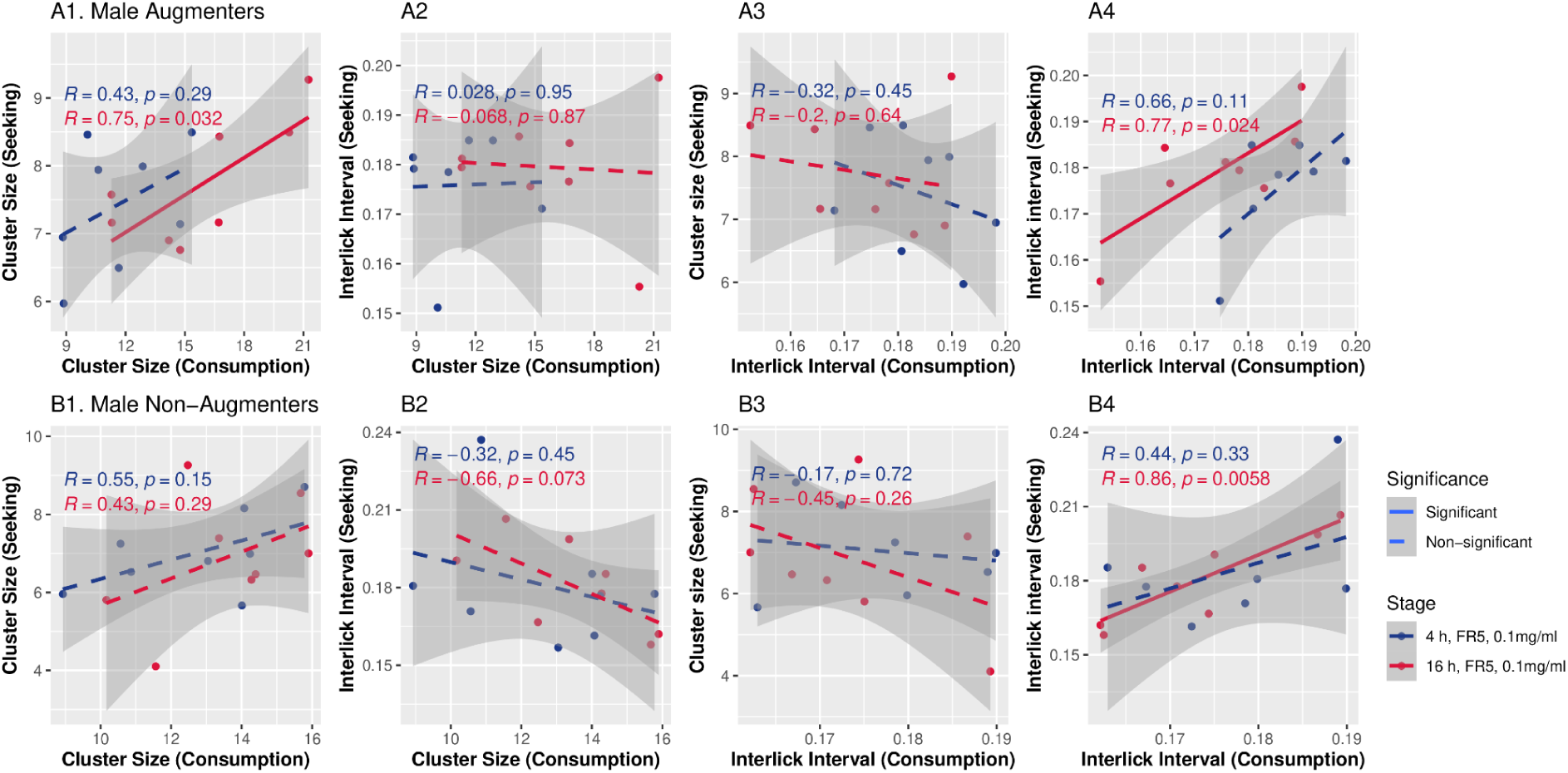
Correlations between lick microstructure parameters during consumption and seeking in male rats. The relationships between CS and ILI during consumption and seeking in male Augmenters (top row, a) and Non-Augmenters (bottom row, b). From left to right, the graphs display the correlation between: (1) consumption CS and seeking CS, (2) consumption CS and seeking ILI, (3) consumption ILI and seeking CS, and (4) consumption ILI and seeking ILI. Each point represents a strain mean. Colors define the 4-hour and 16-hour sessions. The best fit line is solid for significant correlations (p < 0.05) and dotted for non-significant correlations.

## Discussion

We phenotyped a large panel of genetically diverse inbred rat strains to discover strain variation in oral oxycodone self-administration. This effort identified two discrete intake trajectories: “Augmenters” and “Non-Augmenters” consumed comparable amounts of oxycodone during 4-h access, but diverged strikingly during extended 16-h sessions. In a major analytical advance, we powered LMA to distinguish discrete “consumption” from “seeking” lick clusters within a single behavioral model system - a distinction not possible with standard operant lever-press paradigms. This analytical approach revealed the behavioral dynamics underlying augmented oxycodone intake and demonstrated how the hedonic impact of drug consumption affects subsequent drug-seeking behavior.

Several limitations of preclinical models of SUD have been recognized. First, conventional operant paradigms, particularly those employing a fixed-ratio 1 schedule, often conflate appetitive (seeking) and consummatory (taking) behavior^11^, which represent distinct behavioral processes^12^ regulated by different neurobiological mechanisms^13^. Attempts to separate these phases often preclude precise measurements of consumption^14,15^. Second, a direct and quantitative measure of hedonic value, or “liking,” during self-administration is usually lacking. The oral operant licking employed herein is attractive because operant licking produces a reward that is immediately consumed. Advanced LMA distinguishes behavior associated with consumption of the current reward from that involved in seeking the next reward. Moreover, leveraging the established interpretation of cluster size, a quantitative measure of hedonic response^7^, enables the study of hedonic “liking” and its contribution to enhanced “wanting” within the same operant model.

The classification of lick clusters into consumption versus seeking (Figure 3D) is supported by converging empirical evidence. First, cluster size exhibits a sharp, monotonic decline across the initial three clusters. This pattern mirrors the temporal dynamics observed for sucrose licking within the first 5 seconds after consumption^16^ and is consistent with sensory adaptation during ingestive behavior^17^. Second, significantly shorter ILIs during consumption reflect the rapid, stereotyped oral-motor patterns of ingestive behavior, while longer ILIs during seeking suggest a less automated, exploratory licking pattern distinct from active consumption^18^. Beyond the second cluster, the stabilization of both ILI and CS marks an inflection around cluster 3, indicating a behavioral transition from reward consumption to seeking the next reward. This temporal dissociation underscores the ability of the operant licking model to distinguish drug-seeking behavior from consummatory actions.

Operant responding during the unrewarded timeout period is often interpreted as compulsive drug-seeking^19,20^. Some animals display high timeout responding independent of drug consumed during basal training^19,20^. These patterns parallel persistent drug-seeking in human substance use disorders, where drug-associated cues sustain responding during prolonged unavailability^21^.

Timeout responding can also be interpreted within an effort-based motivational framework. In progressive ratio schedules, motivation is quantified by the *breakpoint*, the maximal effort exerted for a single reward^22^. Similarly, behavioral economics methods quantify reward value by decreasing consumption as the effort to obtain reward increases^23^. By analogy, responding during the non-rewarded interval reflects instrumental effort, with lick cluster frequency measuring motivation to obtain the next reward. Compulsive and motivational perspectives thus capture complementary aspects of the same process, where a high level of persistent effort in the absence of reward may represent dysregulated motivation that underlies compulsive seeking.

We found dysregulation of drug-liking and drug-wanting that underlies the vulnerability to increased oxycodone intake in Augmenters. A surge in the frequency of seeking and consumption clusters emitted by Augmenters emerged at the 16-h stage (Fig. 4), paralleling the pattern observed for total oxycodone intake (Fig. 1B, 1C). During this extended access, Augmenters in both sexes emitted significantly more seeking clusters, reflecting heightened motivation to obtain drug, and significantly more consumption clusters, indicating more frequent drug-taking episodes.

While this coordinated change in the regulation of drug taking and seeking could theoretically stem from a shared deficit, such as impaired inhibitory control^24^ or altered satiety mechanisms^25^, a more parsimonious explanation aligns with incentive-sensitization theory^3^ that posits enhanced motivation as the primary driver. This is strongly supported by a novel finding: during 16-h sessions, a robust correlation emerged between consumption cluster size and seeking cluster size in Augmenters of both sexes, a relationship absent in Non-Augmenters (Figure 6A1 and 7A1). This suggests that a heightened hedonic response (“liking”) during consumption elevates the “expected liking” of the drug anticipated during seeking, which subsequently drives the increase in seeking clusters emitted (“wanting”).

This coordinated dysregulation of liking and wanting implicates several neural mechanisms. Candidate pathways include dysregulated mesolimbic dopamine signaling^26^ , which governs both motivation and consummatory actions. This likely involves specific downstream molecular pathways, including nucleus accumbens cAMP-dependent signaling, which is critical for sustained responding requiring high-effort^27^. Additional mechanisms, such as altered mu-opioid receptor function^28^ may also contribute.

In female Augmenters, the hedonic response to oxycodone is tightly coupled to drug seeking. Motivation to seek greater amounts of oxycodone emerges during extended access (Figure 4), yet increases in intrinsic hedonic value, evident in larger consumption CS (Figure 5) and shorter ILI (Supplementary Figure 1), appear as early as 4-h sessions. During extended access, larger consumption CS and shorter ILI not only predict corresponding changes in seeking CS and ILI, but these two parameters are also strongly correlated (Figure 6A), suggesting a reorganization of motor patterns that allows the positive hedonic experience of consumption to directly invigorate subsequent seeking. This finding provides a potential behavioral signature for the vulnerability to augmented oxycodone intake, and suggests an extension to the original incentive-sensitization theory, which posits a dissociation between “wanting” and “liking“^3^. Our findings indicate that these two processes can become tightly intertwined, creating a powerful feedback loop where the hedonic experience of the drug directly fuels the motivation to seek more. This integration of “liking” and “wanting” may drive the transition to compulsive drug use in certain subgroups.

Strong sex differences were evident. Similar to other studies, females consumed more oxycodone than males^29^. In male Augmenters, the hedonic impact of oxycodone intake on subsequent seeking was more compartmentalized. For example, consumption CS strongly correlated with seeking CS during extended access (as in females), but not seeking ILI (Figure 7). This contrasts sharply with female Augmenters, where cross-parameter correlations were strong. Despite the weaker hedonic link between consumption and seeking in male Augmenters, the tendency toward greater hedonic value in Augmenters during consumption at 16-h (Figure 5, consumption CS, p=0.06) may still contribute to the significantly increased number of seeking clusters emitted (Figure 4) - functioning as a common mechanism driving augmentation in both sexes.

Several limitations are acknowledged. First, the classification criteria, based on delivering 60 μL solution (fixed-ratio 5) followed by a 20-second timeout, may need adjustments in other experimental settings. Second, oral self-administration involves pharmacokinetic parameters and potential taste confounds that differ from IVSA, although the consistent licking patterns suggest reinforcing efficacy overcame any potential aversion. Finally, multiple inbred strains provide genetic diversity, yet the specific genetic factors driving the Augmenter phenotype remain unidentified and generalization to outbred populations requires confirmation.

In conclusion, our advanced LMA of lick clusters provides a novel method for dissecting the motivational and hedonic components of drug intake. Strain-dependent Augmenter and Non-Augmenter oxycodone intake phenotypes were differentiated by the frequency of oxycodone lick clusters during consumption and seeking throughout extended drug access and by the increased hedonic response to the drug, which was associated with increased drug seeking in female Augmenters even before intake varied between the two phenotypes. Therefore, the hedonic value of oxycodone during consumption is a powerful strain-dependent enhancer of “wanting” and oxycodone seeking behavior that drives increased drug consumption, particularly in vulnerable female Augmenters.

## Methods

### Animals

Breeders from the HRDP were obtained from Dr. Melinda R. Dwinell at the Medical College of Wisconsin. All rats were bred on campus and housed in groups in a room with a 12:12 h reversed light cycle (lights off: 9AM-9PM) at the University of Tennessee Health Science Center. Experiments were conducted during the dark phase of this cycle. Each rat, including breeders and offspring, had a radio frequency identification (RFID) tag implanted under its skin for identification purposes. Adult rats (PND 65-90) of both sexes were used in oxycodone self-administration. The Animal Care and Use Committee of The University of Tennessee Health Science Center approved all procedures, which complied with NIH Guidelines for the Care and Use of Laboratory Animals. Animals were sacrificed by thoracotomy under deep isoflurane anesthesia.

### Drugs

Oxycodone HCl, a kind gift by Noramco (Wilmington, DE), was dissolved in distilled water.

### Oral operant oxycodone self-administration

We employed the same methodology as previously reported^10,30^. The operant chamber (Med Associates) featured two lickometers: licks on the active spout following a fixed-ratio 5 (FR5) schedule triggered the immediate release of a 60 μl oxycodone solution (0.025–0.10 mg/ml) onto the spout’s tip, along with the activation of an LED visual cue. A 20-second timeout followed the drug delivery, during which licks on the active spout and any licks on the inactive spout were recorded but had no programmed consequences. Rats had free access to food and water and were kept under a reversed light cycle, being tested in their dark phase.

Training commenced with three daily 1-h FR5 sessions at 0.025 mg/ml oxycodone concentration. Subsequent sessions extended to 4-h and occurred every other day. From session four onwards, we doubled the dose every two sessions up to the maximum dose of 0.1 mg/ml. Rats underwent six sessions at this highest dose, followed by a progressive ratio test on session fourteen. During the progressive ratio (PR) test, the number of licks to obtain a subsequent reward was determined using the exponential formula 5e0.2 × injections − 5, such that the required responses per injection were as follows: 1, 2, 4, 9, 12, 15, 20, 25, 32, 40, 50, etc^31^. The PR sessions ended after 20 min of inactivity. The final ratio completed is the breakpoint. Session lengths then increased to 16-h (4 PM - 8 AM) for three sessions. Extinction sessions were conducted for 1-h daily, without programmed consequences, until licks on the active spout decreased to less than fifty for two consecutive sessions, when, in general, the number of licks on the active vs inactive sprouts were no longer significantly different. A final reinstatement session was carried out where active spout licks triggered only visual cue delivery without oxycodone. Final reinstatement data were not collected on a few strains due to campus closures from unexpected weather conditions. We recorded both the number and timing of oxycodone deliveries as well as licks on active and inactive spouts. Full strain names and sexes of rats involved in oxycodone self-administration, along with the number of rats per strain (minimum is 3) and sex are listed in Supplementary Table 1.

**Table S1.** Sample size of oxycodone oral self-administration data by strain.

| Strain | F | M |
| --- | --- | --- |
| ACI/EurMcwi | 9 | 6 |
| BDIX/NemOda | 5 | 5 |
| BN-Lx/Cub | 7 | 7 |
| BN/NHsdMcwi | 16 | 8 |
| BUF/Mna | 3 | 2 |
| BXH12/Cub | 5 | 8 |
| BXH13/Cub | 6 | 3 |
| BXH2 | 9 | 7 |
| BXH6/Cub | 10 | 5 |
| F344/DuCrI | 6 | 4 |
| F344/NHsd | 6 | NA |
| F344/Stm | 8 | 7 |
| FHH/EurMcwi | 7 | 6 |
| FXLE12/Stm | 11 | 13 |
| FXLE13/Stm | 2 | 4 |
| FXLE14/Stm | 3 | 6 |
| FXLE15/Stm | 7 | 8 |
| FXLE17/Stm | 15 | 9 |
| FXLE19/Stm | 5 | 7 |
| HXB1 | 8 | 3 |
| HXB10 | 10 | 10 |
| HXB10XWMI | 7 | 7 |
| HXB13 | 3 | 5 |
| HXB17 | 5 | 4 |
| HXB18 | 5 | 5 |
| HXB2 | 5 | 7 |
| HXB23 | 5 | 6 |
| HXB31 | 11 | 9 |
| HXB4 | 1 | 8 |
| HXB7 | 4 | 3 |
| LE/Stm | 5 | 4 |
| Lew/NHsd | 8 | 8 |
| LEXF2B/Stm | 5 | 5 |
| LEXF5/Stm | 4 | 5 |
| M520/NMcwi | 9 | 5 |
| MNS/N | 13 | 6 |
| MR/N | 5 | 4 |
| MWF | 5 | 6 |
| SHR/NCrl | 6 | NA |
| SHR/Olalpcv | 8 | 6 |
| SHR/OlalpcvxB<br>n-Lx/Cub | 6 | 6 |
| SHR/OlalpcvxB<br>N/NHsdMcwi | 4 | 2 |
| SR/JrHsd | 6 | 2 |
| SS/JrHsdMcwi | 8 | 12 |
| SS/JrHsdMcwi<br>Crl | 9 | 4 |
| SS/JrHsdMcwix<br>SHR/Olalpcv | 9 | 7 |
| WAG/RijCrl | 6 | 7 |
| WLI/Eer | 6 | 3 |
| WMI/Eer | 6 | 5 |

### Identification of Augmenter vs Non-Augmenter Strains

Oxycodone intake at each self-administration stage was calculated separately by sex. The median fold increase in intake from the 0.1 mg/ml 4-h to 16-h stages was approximately 2.5-fold for both sexes. Augmenter strains were identified as those with 16-h intake above the median of each sex and a fold increase above 2.5-fold. This yielded 7 female augmenter strains and 8 male augmenter strains. The female augmenter strains are BDIX/NemOda, FXLE15/Stm, WAG/RijCrl, LEXF2B/Stm, HXB4/Ipcv, BXH12/Cub, and FXLE19/Stm. The male augmenter strains are LE/Stm, BDIX/NemOda, LEXF5/Stm, BN/NHsdMcwi, F344/Stm, FXLE15/Stm, FXLE14/Stm, and BXH2/Cub. For each augmenter strain, one strain that ranked immediately above or below it at the 4-h stage and was not already identified as an augmenter or non-augmenter was designated as the control non-augmenter strain. The female control strains are HXB23/Ipcv, SHR/Olalpcv x Bn-Lx/Cub F1, HXB17/Ipcv, BXH2/Cub, BXH13/Cub, BN-Lx/Cub, and HXB1/Ipcv. The male control strains are FXLE17/Stm, LN/MavRrrc, F344/DuCrl, FXLE13/Stm, LL/MavRrrc, MNS/N, SHR/Olalpcv x Bn-Lx/Cub F1, and HXB2/Ipcv.

### Lick microstructure analysis

Lick microstructure analysis was performed on lick timing data collected from active and inactive spouts, along with corresponding reward delivery times extracted from MedPC data files. Custom R scripts (R Version 4.4.3)^32^ utilizing the dplyr^33^ and lubridate^34^ packages were employed for this analysis. Initial versions of the lick microstructure function were generated with the assistance of Google Gemini 2.5 Pro Experimental (released on 2025-03-25). The output of all parameters from the microstructure function were manually validated using a small synthetic dataset and a few real datasets before the function was used to analyze the entire dataset. Lick clusters were identified based on temporal proximity. A cluster was initiated by a lick, and subsequent licks were included if the ILI did not exceed a pre-defined threshold of 0.5 seconds ^35,36^. A cluster concluded when this ILI threshold was surpassed or when no further licks occurred.

Only clusters comprising at least three licks were included in subsequent analyses. For each valid cluster, several characteristics were quantified: the total number of licks within the cluster (CS), the average ILI between consecutive licks within that cluster, and its functional classification based on reward proximity. Clusters were designated as ‘Overlap’ if a reward delivery occurred within the cluster’s temporal boundaries, or ‘No_Overlap’ otherwise. To examine licking patterns specifically following a reward, ‘No_Overlap’ clusters were assigned a sequential run index. This index started at 1 for the first ‘No_Overlap’ cluster after a reward delivery (or after the session start if no prior reward had occurred) and incremented for each subsequent consecutive ‘No_Overlap’ cluster. This count was reset to zero after the 20-second post-reward timeout interval when an ‘Overlap’ cluster occurred or if a reward was delivered between clusters. For analytical consistency, all ‘Overlap’ clusters were assigned a run index of 0. Latency following reward is the time between a reward delivery and the first lick of each licking cluster that follows that reward. Each cluster is assigned to the reward immediately preceding it, including clusters that occur during the timeout and those that occur afterward, up until the next reward. All clusters initiated during the 20-second post-reward timeout period following a reward were specifically analyzed to determine multiple parameters (see next section) defining the parameters for lick function classification.

### Functional classification of lick clusters

Licking behavior was subsequently categorized into distinct functional categories based on the cluster type and its sequential run index following a reward. ‘Consumption’ was defined as licking behavior primarily associated with reinforcer intake, encompassing all ‘Overlap’ clusters (which had a run index of 0) and ‘No_Overlap’ clusters that were the first or second in sequence after a reward. ‘Seeking’ was defined as licking behavior not immediately tied to consumption of a reward, consisting of ‘No_Overlap’ clusters that were third or later in the sequence following a reward. This classification scheme was informed by preliminary mixed-effects model analyses of cluster characteristics (e.g., CS, ILI) against the sequential run index, which revealed distinct properties for these initial post-reward clusters compared to later ones.

### Statistical analysis

Outliers were addressed by first applying a logarithmic transformation to each variable to normalize distributions. Outliers were then identified on this transformed scale as values falling below the first quartile minus twice the interquartile range (IQR) or above the third quartile plus twice the IQR. These identified outliers were treated as missing data, and the remaining data were subsequently back-transformed to their original scale. To prepare data for statistical comparisons between Augmenter and Non-Augmenter intake phenotypes, the detailed cluster data, following outlier removal and lick function categorization, underwent a two-stage aggregation process. First, daily mean values for the number of clusters, total licks, average CS, and average intra-cluster ILI were computed for each individual animal, within each experimental stage, sex, and assigned intake phenotype. These calculations were performed separately for clusters categorized as ‘Consumption’ and ‘Seeking’. Second, these daily animal means were further averaged across all experimental days to yield a single mean value per strain for each combination of sex, intake phenotype, experimental stage, and lick function. These strain-level means served as the primary data points for subsequent statistical modeling.

An estimate of narrow-sense heritability (i.e. the proportion of total phenotypic variation that is due to the *additive* effects of genes (h^2^) for nicotine or food reward was obtained using the formula: *h*^2^ *=V_A_/(V_A_+V_E_)*^37,38^

All statistical analyses were performed using R (Version 4.4.3). Linear mixed-effects models (LMMs) were implemented using the lme4^39^ and lmerTest^40^ packages, with post-hoc comparisons conducted via the emmeans^41^ package. The dependent variables, derived from the strain-level aggregated data, included the number of clusters, total licks, mean CS, and mean intra-cluster ILI. The LMMs incorporated experimental stage, sex, intake phenotype, and lick function as fixed effects, along with all their potential interactions. To account for inherent variability due to genetic background, ‘strain’ was included as a random intercept in these models. The significance of fixed effects and their interactions was assessed using Type III Analysis of Variance (ANOVA) tables, with Satterthwaite’s method for approximating degrees of freedom. Following significant main effects or interactions, post-hoc pairwise comparisons of estimated marginal means were performed, applying Tukey’s HSD method for multiple comparison adjustment. A significance level of p < 0.05 was adopted for all statistical inferences.

## Acknowledgments

The authors would like to express their gratitude to Dr. Melinda R. Dwinell (Medical College of Wisconsin) for generously providing the breeders for the Hybrid Rat Diversity Panel (HRDP), and to the Center for Integrative and Translational Genomics at the University of Tennessee Health Science Center for its support in maintaining the HRDP. We also thank Ms. Caroline Jones and Ms. Anna Fracchia for maintaining the breeding colony, and Dr. Shuangying Leng for conducting the oral self-administration experiments. This study was supported by NIDA U01DA053672 (B.M. Sharp, R.W. Williams and H. Chen).

